# When, why and how clonal diversity predicts future tumour growth

**DOI:** 10.1101/2019.12.17.879270

**Authors:** Robert Noble, John T Burley, Cécile Le Sueur, Michael E Hochberg

**Author notes:** RN and JTB should be considered joint first author.

## Abstract

Intratumour heterogeneity holds promise as a prognostic biomarker in multiple cancer types. However, the relationship between this marker and its clinical impact is mediated by an evolutionary process that is not well understood. Here we employ a spatial computational model of tumour evolution to assess when, why and how intratumour heterogeneity can be used to forecast tumour growth rate, an important predictor of clinical progression. We identify three conditions that can lead to a positive correlation between clonal diversity and subsequent growth rate: diversity is measured early in tumour development; selective sweeps are rare; and/or tumours vary in the rate at which they acquire driver mutations. Opposite conditions typically lead to negative correlation. Our results further suggest that prognosis can be better predicted on the basis of both clonal diversity and genomic instability than either factor alone. Nevertheless, we find that, for predicting tumour growth, clonal diversity is likely to perform worse than conventional measures of tumour stage and grade. We thus offer explanations – grounded in evolutionary theory – for empirical findings in various cancers. Our work informs the search for new prognostic biomarkers and contributes to the development of predictive oncology.

## Introduction

Despite a wealth of available genomic data, the development of reliable predictive tools in oncology remains challenging (Yankeelov et al., 2015; Lipinski et al., 2016; Turajlic et al., 2019). Although the search for prognostic biomarkers has largely focussed on the presence or absence of particular genomic aberrations, indices of intratumour heterogeneity, which depend only on aberration frequencies, not identities, have emerged as promising alternative predictors (Marusyk et al., 2012; Polyak, 2014; Alizadeh et al., 2015; Jamal-Hanjani et al., 2015; Maley et al., 2017). Higher clonal diversity has been found to predict worse clinical outcome in Barrett’s oesophagus (Maley et al., 2006; Merlo 2010; Martinez et al., 2016), ovarian cancer (Schwarz et al., 2015), lung cancer (Jamal-Hanjani et al., 2017) and breast cancer (Park et al., 2010; Rye et al., 2018). Computational modelling of pre-malignant somatic evolution further indicates that genetic diversity indices can predict cancer risk more reliably than the presence or absence of particular mutations (Dhawan et al., 2016). However, studies in kidney cancer (Turajlic et al., 2018) and across cancer types (Andor et al., 2015) have found more complicated relationships between intratumour heterogeneity and its clinical impact, which elude simple explanation. It also remains unclear whether and how intratumour heterogeneity can complement established prognostic biomarkers – such as tumour stage and grade – and other proposed ecological and evolutionary indices (Maley et al., 2017).

Interpreting intratumour heterogeneity is challenging due to the complexity of tumour evolutionary dynamics (Gerlinger et al., 2012; Burrell et al., 2013; Lipinski et al., 2016; Cross et al., 2016; Venkatesan & Swanton, 2016; Williams et al., 2016). Greater heterogeneity at the scale of tumour clonal composition may reflect higher genomic instability (Hanahan & Weinberg, 2011) and may correspond to a greater likelihood of malignant cell phenotypes being present (Maley et al., 2006; Greaves & Maley, 2012; Maley et al., 2017). On the other hand, clonal sweeps initiated by highly adapted clones might reduce diversity within aggressive tumours (Maley et al., 2006; Robertson-Tessi & Anderson, 2015; Maley et al., 2017). The frequency of such selective sweeps in turn depends on the extent of clonal interference (Lang et al., 2013; Martens et al., 2011) and spatial constraints (Michor et al., 2003; Noble et al., 2019), which may vary between tumour types. Potentially, clonal diversity could fail as a prognostic biomarker either because it is insufficiently variable within patient cohorts or because tumour progression is so stochastic as to be effectively unpredictable. Even when forecasts are possible, their accuracy is expected to depend on the number and location of cells used to assess heterogeneity. Some tumour regions will tend to harbour more clonal diversity than others, and a clone’s ability to expand and contribute to growth rate is constrained by intracellular interactions, which depend on the clone’s location relative to other clones and to the tumour edge.

Here we use a computational model of solid tumour evolution to characterize the relationship between clonal diversity and subsequent tumour growth rate, an important factor in determining morbidity and mortality risk. We identify conditions that determine the sign and strength of correlations between these two variables. We thus provide insight into the evolutionary processes underlying clinical observations. Our results contribute to establishing a theoretical foundation for predictive oncology.

## Results

### Overview of computational model and analysis

We simulate invasive tumour growth and evolution using a spatial, stochastic computational model specifically designed to recapitulate the spatial structure of common acinar tumour types, such as kidney, lung and breast carcinomas. This model has previously been shown to generate a pattern of branched evolution, consistent with previous observations in various cancer types (Noble et al., 2019). Our model represents tissue as a two-dimensional regular grid of “demes”, corresponding to localised subpopulations of interacting cells. Initially, all demes contain normal cells, except that one deme at the centre contains a single tumour cell with a higher division rate than normal cells. Tumour cells stochastically divide, mutate, die, and disperse between neighbouring demes, whereas normal cells undergo stochastic division and death only. The number of cells per deme is regulated, via negative feedback, to remain approximately constant. Specifically, we assume that the cell death rate increases from zero to a large value (100 times the initial cell division rate) when the deme population size exceeds carrying capacity *K*. Whenever a tumour cell is created via cell division, it either remains in the same deme or, with probability *m*, disperses to a neighbouring deme. The value of dispersal probability *m* is adjusted for the value of *K*, such that, in the absence of mutation, all tumours would take a similar amount of time to reach the endpoint size (corresponding to several years in real time). We restrict our analysis to mutations that increase tumour cell fitness, termed driver mutations. Driver mutations occur at rate *μ* and individually increase cell division rate *r* by a factor of 1 + *X*(1 – *r* / *m*), where *X* is drawn from an exponential distribution with mean value *s*, and *m* is an upper bound on the cell division rate. Because we set *m* to be much larger than the initial value of *r*, the combined effect of drivers is approximately multiplicative. Canonical parameter values are listed in Supplementary table 1.

We focus our study on how the principal parameters of our model – deme carrying capacity (*K*), driver mutation rate (*μ*) and mean driver fitness effect (*s*) – influence the coefficient of correlation between a predictor variable (such as clonal diversity), measured at a particular tumour size, and the rate at which the tumour grows to a larger endpoint size (Figure 1a). We define the tumour’s future growth rate as

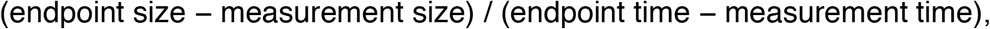

where measurement size is the tumour size when the predictor variable is measured; measurement time is the corresponding tumour age; and endpoint size and endpoint time are the tumour size and age, respectively, when the tumour reaches the endpoint size. We use Spearman’s rank correlation coefficient so as to allow for non-linear correlations. Correlation coefficients closer to 1 or −1 correspond to more reliable forecasts.

**Figure 1.**
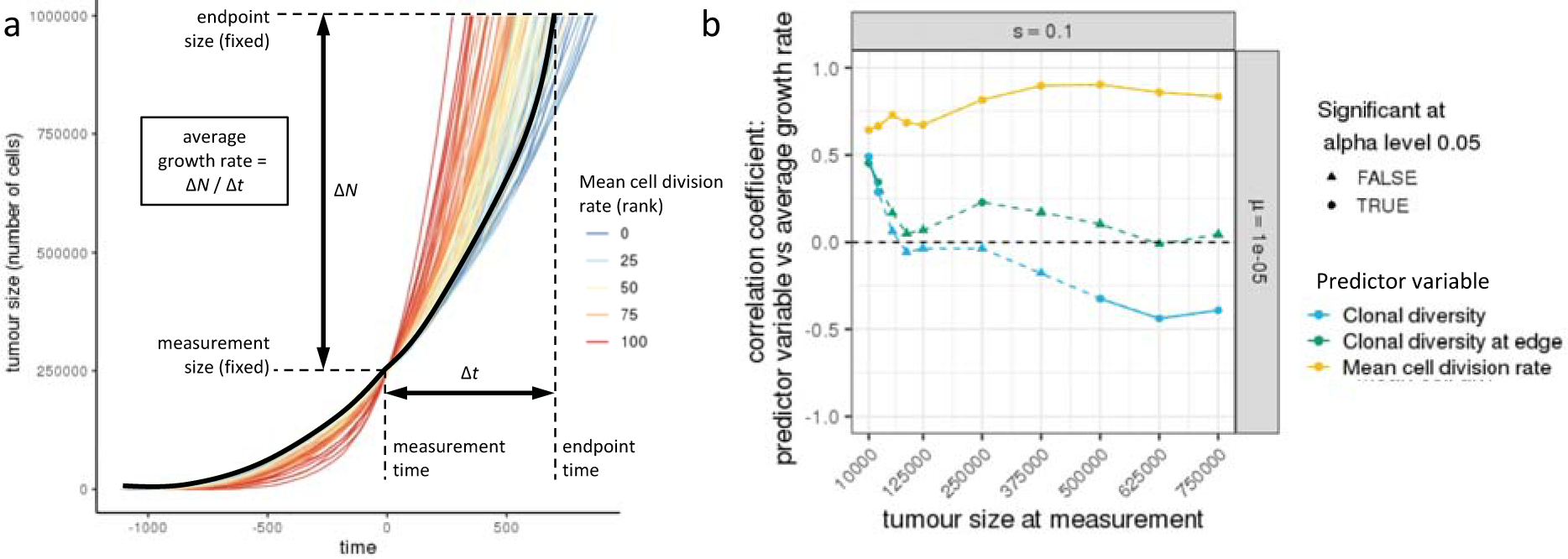
Correlations between predictor variables and future tumour growth rate for cohorts of tumours with identical parameter values. a) Example tumour growth trajectories. In this cohort, tumours with higher mean cell division rate (red curves) typically have higher future growth rate, which results in a positive correlation coefficient. One trajectory is shown in black to illustrate the method used to calculate average growth rate. b) Correlation coefficients between predictor variables and subsequent tumour growth rate, for different measurement sizes. Correlation coefficients that significantly differ from zero (p < 0.05) are indicated by circular points and solid lines, as opposed to triangular points and dashed lines. Each point represents a cohort of 100 simulated tumours; lines are added only to guide the eye. Parameter values are *K* = 512, *μ* = 10^−5^, *s* = 0.1.

### Mean cell division rate reliably predicts future tumour growth

Before considering clonal diversity, we first look at the predictive value of mean cell division rate in our model. Cell division rate is an important component of conventional clinical grading of tumours and therefore provides a benchmark with which we can compare alternative predictor variables. To test the limits of forecastability, we examine cohorts of simulated tumours with identical parameter values and initial conditions, so that variation among growing, evolving tumours is entirely due to stochastic events.

In line with intuition, and consistent with the clinical evidence that underpins cancer staging, mean cell division rate is consistently positively correlated with future tumour growth rate in our model (Figure 1b, yellow curve; Supplementary figure 1). The correlation is however imperfect because tumour growth trajectories continue to be swayed by the stochastic accumulation of driver mutations between the measurement time and the endpoint time (the time at which the tumour reaches the endpoint size, as shown in Figure 1a). Correlations are weaker when cell division rate is measured at a very early stage of tumour growth, especially when driver mutations have only small effects on cell fitness (Supplementary figure 1). This is mostly because very small tumours typically lack sub-clonal driver mutations and hence there is insufficient variation in cell division rate to support a correlation. When driver mutations have large fitness effects, correlations are weaker if cell division rate is measured at a very late stage (Supplementary figure 1). In general, these initial results demonstrate that, despite the effects of ongoing stochastic events, tumour growth in our models is in principle forecastable.

### Diversity-growth correlation depends on when and where diversity is measured

Having established potential for forecasting, we next examine correlations between clonal diversity and subsequent tumour growth rate, which we will refer to as diversity-growth correlations. We define diversity in terms of Simpson’s index (Methods) and define a clone as a set of tumour cells with the same combination of driver mutations (that is, mutations that increase cell fitness). This definition of a clone is consistent with that used by the TRACERx Renal Consortium, which has conducted the most sophisticated clinical investigation of the association between intratumour heterogeneity and disease progression to date (Turajlic et al., 2018). When measured at an early stage of tumour development, we find that clonal diversity is positively correlated with subsequent tumour growth rate. Conversely, clonal diversity measured at a later stage can be negatively correlated with tumour growth rate (Figure 1b, blue curve). Therefore diversity-growth correlations importantly depend on the stage of tumour development when diversity is measured.

One factor that affects the predictive value of clonal diversity is that cells contribute unequally to tumour growth. Since our model assumes that tumours grow by invading normal tissue, the build-up of driver mutations in the tumour core does not influence total population growth unless an especially fit clone evades clonal interference and spreads so rapidly through the tumour so as to overtake the expanding edge. As such we would predict that adaptive mutations arising near the tumour edge are more likely to increase tumour growth rate, at least over short timescales. Consistent with these considerations, negative diversity-growth correlations are absent or less pronounced when diversity is measured only in cells sampled from the tumour edge (Figure 1b, green curve; Supplementary figure 1).

### Diversity-growth correlation depends on the extent of clonal turnover

A potential explanation for negative diversity-growth correlation, typically observed when clonal diversity is measured at larger tumour sizes (Figure 1b), is the occurrence of selective sweeps that purge heterogeneity. To test this hypothesis, we use a clonal turnover index (Methods) to quantify changes in clone frequencies over time. As expected, clonal diversity, when measured at an intermediate tumour size, is positively correlated with future tumour growth rate only in cohorts that have low clonal turnover, whereas correlations are negative in cohorts that have high clonal turnover (Figure 2a). Low clonal turnover corresponds to one or more of three factors: low driver mutation rate, low driver fitness effect, and/or small deme carrying capacity (Supplementary figure 2).

**Figure 2.**
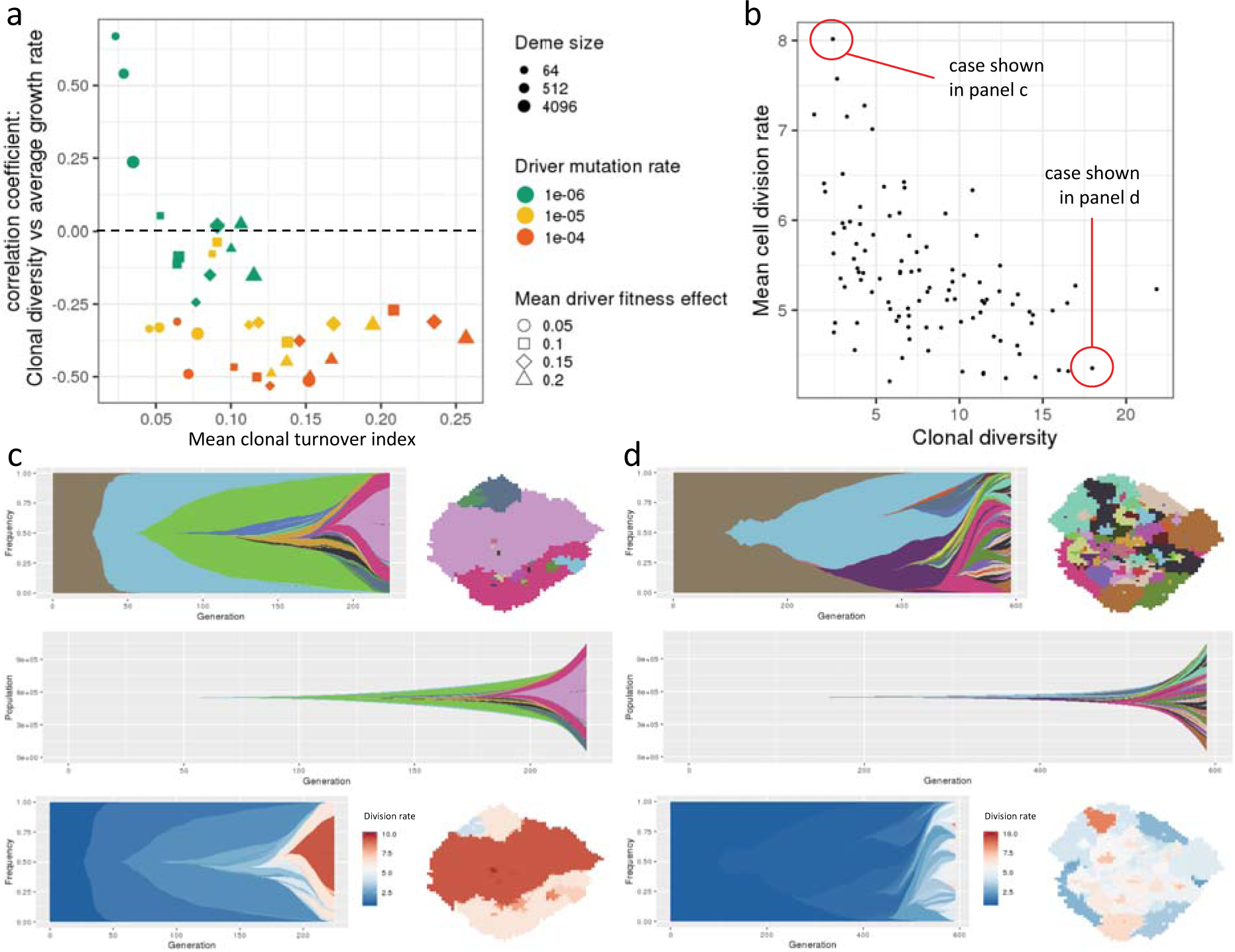
Higher clonal diversity can be associated with slower future tumour growth. a) Correlation coefficients between clonal diversity (measured at a tumour size of 250,000 cells) and subsequent tumour growth rate, plotted against mean clonal turnover index. Each point represents a cohort of 100 simulated tumours. b) An example of a tumour cohort exhibiting negative correlation between clonal diversity (measured at a tumour size of 750,000 cells) and mean cell division rate (Spearman’s correlation coefficient −0.55; *p* < 10^−8^). c) A simulated tumour from this cohort exhibiting a succession of selective sweeps, resulting in high mean cell division rate, high growth rate and low clonal diversity. Top left is a Muller plot in which colours represent clones with distinct combinations of driver mutations (the original clone is grey-brown; subsequent clones are coloured using a recycled palette of 26 colours). Descendant clones are shown emerging from inside their parents. Top right is a spatial plot of the tumour at the endpoint time, in which each pixel corresponds to a deme containing approximately *K* cells, coloured according to the most abundant clone within the deme. The middle row shows a Muller plot of clone sizes, rather than frequencies. In the bottom row, clones in the Muller and spatial plots are coloured by cell division rate. d) A simulated tumour from the same cohort exhibiting low mean cell division rate, low growth rate and high clonal diversity due to extensive clonal interference. Parameter values in panels b-d are *K* = 512, *μ* = 10^−5^, *s* = 0.2.

In cohorts with high clonal turnover, we observe very similar negative correlations between clonal diversity and contemporaneous mean cell division rate (Figure 2b; Suppmentary figure 3). Examining the evolutionary dynamics of individual tumours within such cohorts, we find that the combination of low clonal diversity, high mean cell division rate and high tumour growth rate is characteristic of a succession of selective sweeps (Figure 2c). Conversely, high clonal diversity, low mean cell division rate and low tumour growth rate result from clonal interference between less well-adapted clones (Figure 2d).

### Clonal diversity is positively correlated with future growth rate among tumours with diverse driver mutation rates

It is reasonable to expect that, in reality, even tumours of the same size and type (morphology and anatomical location) vary in their underlying biological parameters because of intrinsic and microenvironmental factors. In particular, genomic instability and mutation burden are highly variable within cancer types (Chalmers et al., 2017) and – at least across cancer types – the number of driver mutations per tumour generally increases with total mutation burden (Martincorena et al., 2017). To examine the consequences of such variation, we next analyse cohorts containing tumours with differing values of the driver mutation rate, *μ*.

In such cohorts, we find that future growth rate is consistently positively correlated with clonal diversity (Figure 3a, blue curves; Supplementary figure 4a). Correlations are moreover stronger when diversity is measured across the entire tumour, rather than only at the tumour edge (Figure 3a, blue versus green curves). In either case, correlation coefficients are insensitive to varying the endpoint size (Figure 3b). These results demonstrate that parameter variability within a cohort can transform diversity-growth correlations, provided that intrinsic, deterministic differences between tumours outweigh the stochastic differences that emerge during evolution. Nevertheless, we find that, as a predictor of future growth rate, clonal diversity is always inferior to mean cell division rate (Figure 3a, yellow curves; Supplementary figure 4a).

**Figure 3.**
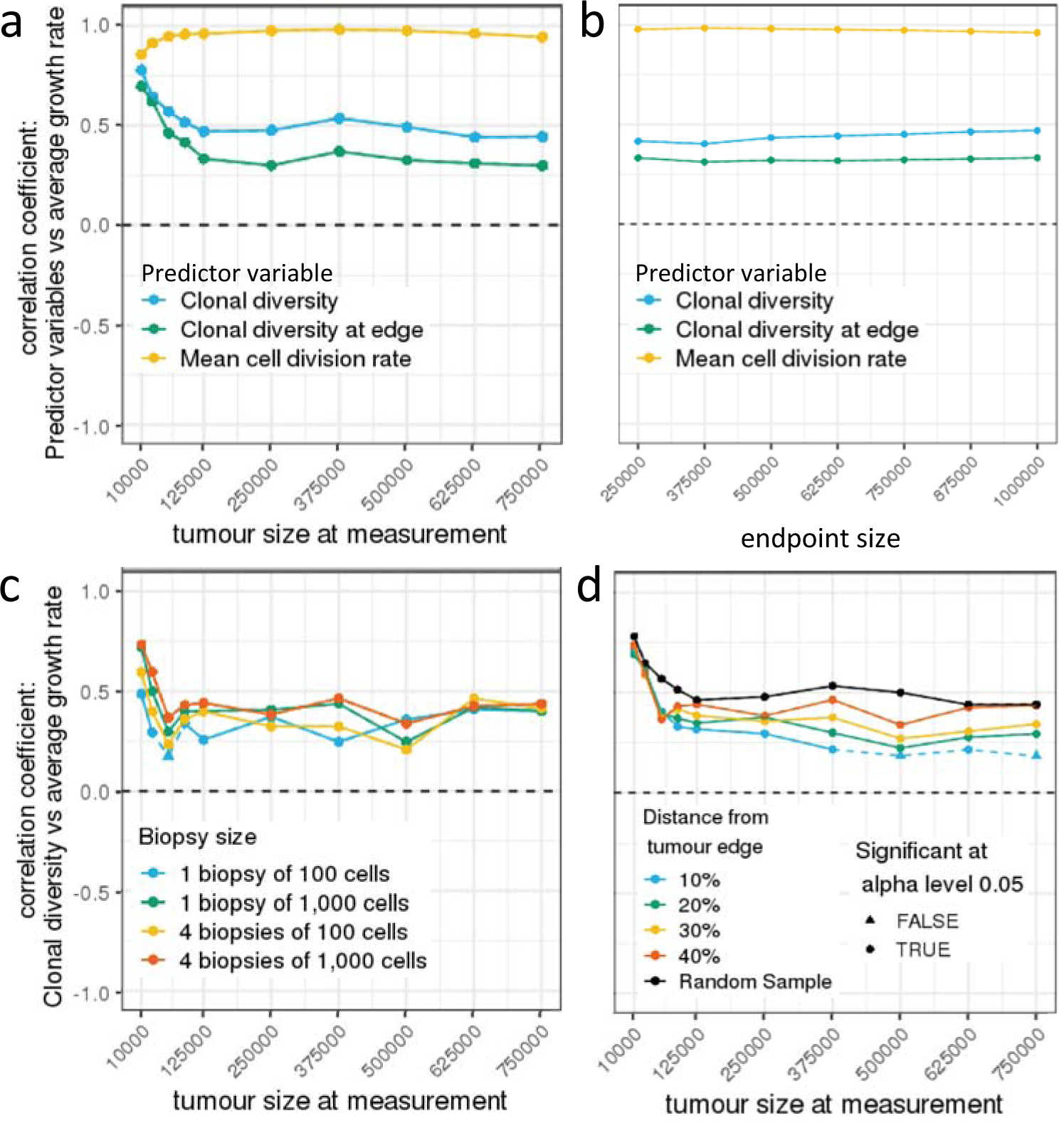
Correlation coefficients for cohorts of tumours with diverse driver mutation rates. a) Correlation coefficients between predictor variables and subsequent tumour growth rate, for different measurement sizes. b) Correlation coefficients between predictor variables (measured at a tumour size of 125,000 cells) and subsequent tumour growth rate, for different endpoint sizes. c) Correlation coefficients between future tumour growth rate and clonal diversity, measured in biopsy samples of different number and size (at a depth of 40% from the tumour edge), for different measurement sizes. d) Correlation coefficients between future tumour growth rate and clonal diversity, measured in biopsy samples taken at different depths relative to the tumour edge (four biopsy samples, each of 1,000 cells, are taken from each tumour), for different measurement sizes. Correlation coefficients that significantly differ from zero (*p* < 0.05) are indicated by circular points and solid lines, as opposed to triangular points and dashed lines. In all plots, each point represents a cohort of 100 simulated tumours; lines are added only to guide the eye. Parameter values are *K* = 512 and *s* = 0.1. The driver mutation rate of each tumour is chosen by sampling a random value *X* from a uniform distribution between 4 and 6 and then setting *μ* = 10^−*X*^.

In cohorts of tumours with differing driver mutation rates, high tumour growth rate correlates not only with high clonal diversity but also with high driver mutation rate. Importantly, however, clonal diversity is not merely a proxy for driver mutation rate. For example, as previously noted, diversity can be positively correlated with subsequent growth rate in tumours with low values of *μ*, and yet negatively correlated with subsequent growth rate in tumours with high values of *μ*. Hence prognosis can be better predicted on the basis of both parameters than either one alone. This finding can be readily observed in Kaplan-Meier plots of progression to endpoint size (Figure 4a).

**Figure 4.**
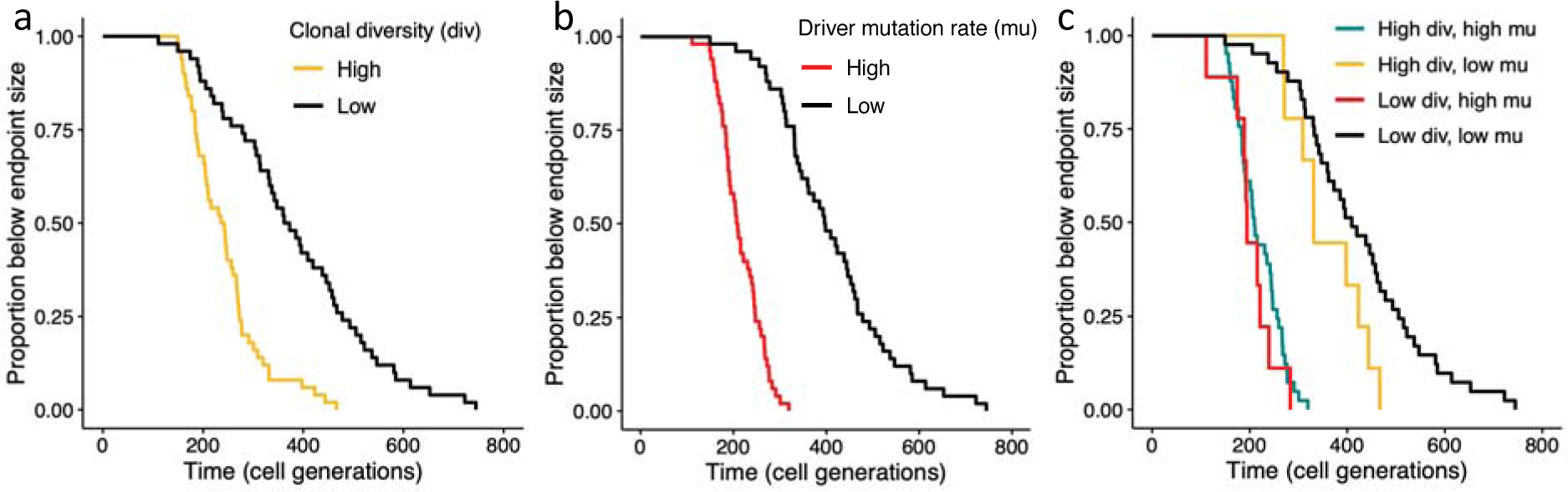
Kaplan-Meier for progression to endpoint tumour size in a single cohort of tumours with diverse driver mutation rates. Tumours are grouped by clonal diversity (a); driver mutation rate (b); or both clonal diversity and driver mutation rate (c). For grouping, values above and below the median are classed as high and low, respectively. Parameter values are *K* = 512 and *s* = 0.05. The measurement size is 250,000 cells and the endpoint size is one million cells. The driver mutation rate of each tumour is chosen by sampling a random value *X* from a uniform distribution between 4 and 6 and then setting *μ* = 10^−*X*^.

When tumours differ in the mean driver fitness effect (parameter *s*) but not in driver mutation rate (*μ*), diversity-growth correlations instead follow the same pattern as when parameter values do not vary (Supplementary figure 5).

### The predictive value of clonal diversity is robust to biopsy sampling bias

Clinical evaluations of intratumour heterogeneity are typically based on examining cells in one or more relatively small biopsy samples. To assess how such sampling bias affects our results, we repeated our analyses using clonal diversity measurements derived only from biopsies (Methods). When driver mutation rate varies within cohorts, results based on biopsies are similar to those based on all tumour cells, except when the sample size is very small (Figure 3c; Supplementary figure 4b). Also when driver mutation rate varies within cohorts, diversity-growth correlations are stronger when biopsy samples are taken from inside the tumour, rather than from the edge (Figure 3d; Supplementary figure 4c).

## Discussion

Intratumour heterogeneity is considered a promising prognostic biomarker because it directly relates to the evolutionary processes that drive tumour progression (Marusyk et al., 2012; Polyak, 2014; Alizadeh et al., 2015; Jamal-Hanjani et al., 2015; Maley et al., 2017). The general nature of this relationship has, however, proven difficult to characterize. By simulating tumour evolution under different conditions in numerous virtual patient cohorts, here we have taken first steps towards disentangling when, why and how clonal diversity (a particular form of intratumour heterogeneity) predicts tumour growth rate, which is in turn an important predictor of clinical progression.

We have identified three mutually-reinforcing factors that can lead to positive correlations between clonal diversity and future tumour growth. First, positive diversity-growth correlations are seen when clonal diversity is measured at an early stage of tumour progression. This is because high diversity in a small tumour indicates the presence of mutations that are under selection, and that will eventually accelerate tumour growth, but that have not yet had time to fix. Second, clonal diversity, even when measured at a later stage, can positively correlate with future growth rate provided that tumour spatial structure and/or evolutionary parameters sufficiently restrict the rate of clonal turnover, which otherwise purges diversity within more aggressive tumours. Finally, positive diversity-growth correlations can arise in cohorts of tumours that have differing rates of acquiring beneficial (driver) mutations – for example due to variation in genomic instability – because higher diversity then correlates with higher driver mutation rate, which in turn predicts faster tumour growth. In the absence of all three of these factors, we find that the correlation between clonal diversity and future tumour growth rate is typically negative.

Although the influence of stochastic factors fundamentally limits predictability in our model – as is the case in real cancers (Lipinski et al., 2015) – we find that tumour growth forecasts can be remarkably reliable. Due to complex processes at multiple spatial and temporal scales, forecasts of weather and of many kinds of ecological dynamics become progressively less accurate as they reach further into the future (Petchey et al., 2015). Tumour growth in our model conforms to a very different pattern, such that forecast accuracy is unchanged or even improves as the projection period lengthens (Figure 3b; Supplementary figure 4b). This is because, in a growing population, early mutations are likely to reach higher frequency and hence be more influential than later mutations (which arise in a larger population, typically further from the tumour edge, and encounter more clonal interference), yet even early mutations have somewhat delayed effects on tumour growth, due to the time required for clonal expansion. A tumour’s long-term growth trajectory is mostly determined at an early stage, even as driver mutations occur increasingly often over time.

In their own study cohort and in the larger TCGA kidney cancer cohort, the TRACERx Renal consortium found that low intratumour heterogeneity correlates with longer progression-free and overall survival times when tumours have low genomic instability, but not when tumours have high genomic instability (Turajlic et al., 2018). Our model generates a similar pattern (Figure 4c) and thus provides an explanation for clinical observations, provided that tumour growth rate during relapse (which determines progression-free survival) correlates with the growth rate of the same tumour had it been left untreated (which is the outcome we forecast). From a clinical perspective, however, the crucial question is whether new prognostic biomarkers can outperform the status quo. The TRACERx Renal consortium found that the predictive power of intratumour heterogeneity and genomic instability remained significant after adjusting for known prognostic variables (including stage and grade) in the TCGA kidney cancer cohort but not in their own smaller cohort of 100 tumours (Turajlic et al., 2018). Consistently, in our model results, forecasting growth rate from clonal diversity is inferior to forecasting based on mean cell division rate, a key factor in assigning cancer grade. Nevertheless, it is important to note that we have not tested here whether clonal diversity can predict other clinically important factors, such as metastatic potential or the likelihood of a drug resistant clone being present at the time of initiating treatment.

Our work contributes to a growing body of work on predicting evolution in complex systems, which is gaining increasing attention because of its potential to provide mechanistic explanations for patterns observed in clinical data and to guide therapeutic interventions (reviewed in Lässig et al., 2017). For example, using a non-spatial computational model and analysis of high-throughput sequencing data, Williams and colleagues (2018) recently inferred the strength of selection in various cancer types and used the results to forecast changes in clonal architecture. Hosseini and colleagues (2019) and Diaz-Uriarte and Vasallo (2019) have instead examined the order of accumulation of driver mutations and found that certain types of fitness landscape lead to more or less predictable tumour evolutionary trajectories. Our approach is both distinct and complementary, in that we seek to describe the stochastic processes of clonal initiation, expansion and interaction in a realistic spatial context, and we focus on generic, clinically relevant predictor variables and outcomes.

We have kept our model simple so as to yield the most general insights. Our work thus has several limitations that motivate further investigation. First, even allowing for the fact that we simulate a two-dimensional slice of a larger three-dimensional tumour, the endpoint size of one million cells is unrealistically small. This discrepancy might be relatively inconsequential, however, given our finding that forecast accuracy is robust to increasing endpoint size. In common with previous computational modelling studies (Bozic et al., 2010; Waclaw et al., 2015), we assume an infinite sites model of evolution. We also assume only very weak diminishing-returns epistasis. We expect that assuming a finite sites model or stronger diminishing-returns epistasis – such as observed in long-term *E. coli* evolution (Wiser et al., 2013) – would tend to make tumour growth even more deterministic and more predictable. Relatedly, we neglect deleterious mutations and hence do not impose any fitness cost of high genomic instability, a factor that might help explain slower growth of especially heterogeneous tumours (Andor et al., 2015). We have not attempted to investigate how forecast accuracy might be affected by frequency-dependent cell-cell interactions or microenvironmental heterogeneity. Nor have we explicitly modelled treatment and the evolution of resistance.

An especially promising direction for future research is in combining clonal diversity measurements with information about evolutionary processes, such as the pervasiveness of selective sweeps in a given tumour type across individuals or in a particular tumour within an individual. Although it is generally infeasible to infer a complete history of clonal turnover, genomic data from even a single timepoint can potentially be used to infer the strength of selection, time since the most recent common ancestor, and rate of demographic expansion (Williams et al., 2018; Alves et al., 2019). Future work should employ multi-variable statistical models that incorporate such information about tumour evolution, ecology and demography (Maley et al., 2017). These methods are likely to outperform the simple regression models we have examined here. Future studies should also evaluate system-specific modifications of our model and our biopsy sampling protocol, using multi-omics data, histopathology image analysis, and cancer staging assays to infer sub-clonal cell proliferation rates and to characterize cell-cell interactions. By laying the foundations for such projects, the current study comprises a step towards the ultimate goal of personalized cancer evolution forecasts, parametrised with patient-specific data.

## Author contributions

MEH and RN conceived the research question and designed the study. RN designed and created the computational modelling framework. JTB, CLS and RN wrote analysis code, ran simulations and analysed output data. RN and JTB wrote the manuscript with contributions from MEH and CLS.

## Acknowledgements

We thank Yannis Michalakis, Dominik Burri, Katharina Jahn and Francesco Marass for helpful comments. The Montpellier Bioinformatics Biodiversity platform aided the initial development of this work. MEH acknowledges support from the McDonnell Foundation (Studying Complex Systems Research Award No. 220020294), ITMO Cancer AVIESAN (Program ‘HetColi’) and the Institut National du Cancer (2014-1-PL-BIO-12-IGR-1). RN acknowledges support from ERC Synergy Grant 609883 and the National Cancer Institute of the National Institutes of Health under Award Number U54CA217376. The content is solely the responsibility of the authors and does not necessarily represent the official views of the National Institutes of Health.

## Methods

### Clonal diversity index

We measure clonal diversity using the inverse Simpson index, defined as one divided by the sum of the squares of all clone frequencies. This index can be straightforwardly interpreted as an effective number of distinct clones. For example, if a tumour contains n equally abundant clones then the diversity index is 1 / (*n* × (1/*n*)^2^) = *n*.

### Clonal turnover index

Our clonal turnover index quantifies the average rate of change in clone frequencies. Specifically, for each time point *t* ≥ *δt*, we calculate *E*(*t*) = ∑_*i*_(*f*_*i*_(*t* − *δt*) − *f*_*i*_(*t*))^2^, where *f*_*i*_(*t*) is the frequency of clone *i* at time *t*, and *δt* is 10% of the total simulation time (that is, the time from when the simulation is initiated with a single cell until the endpoint time). The clonal turnover index is then the mean value of *E*(*t*). Time is measured in cell generations (that is, relative to the expected cell division time of the initial tumour clone) and the time points are evenly spaced.

### Sampling

We variously measure clonal diversity among all cells in the simulated tumour, all cells at the tumour edge, a small random sample of cells, or one or more small localised samples of cells (simulated biopsy). We sample the edge of the tumour by selecting all demes that are directly “visible” from one of the four sides of the square lattice and, from each of these edge demes, we sample a number of cells equal to the square root of the deme carrying capacity, *K* (so the total sample size is not biased by deme size).

Biopsies are taken from small disc-shaped regions, centred at a given proportion of the distance from the tumour’s edge to its centre of mass. When four biopsies are taken from the same tumour, they are located on perpendicular transects through the tumour’s centre of mass (Supplementary figure 6). The number of cells sampled from a deme is proportional to the extent of overlap between the biopsy region and the deme. If the sample size is less than the deme population size then we employ multinomial sampling of clones. Whereas we assume a relatively small number of cells per biopsy, it is important to note that our simulation data is free of measurement error. The information obtained from a small biopsy sample without error is comparable to what could be obtained from analysing a larger data set with noise.

### Parameter values

Although cancer genomic data is broadly consistent with driver mutation rates between 10^−6^ and 10^−2^ per cell division (Bozic et al., 2019), most studies have settled on values close to 10^−5^ per cell division (Bozic et al., 2010; Waclaw et al., 2015; Sun et al., 2017). Likewise, we consider values of *μ* in the range 10^−6^–10^−4^.

In certain non-spatial mathematical models, driver fitness effects of less than 1% are consistent with the acquisition of multiple driver mutations during tumour growth (Bozic et al., 2010). Importantly, the same result does not necessarily hold when spatial structure and/or clonal interference inhibit the spread of beneficial mutations. Consistent with recent genomic data analysis (Williams et al., 2018), we assume the mean driver fitness effect *s* to be between 0.05 and 0.2.

We have previously found that glands of invasive, acinar tumours typically contain between a few hundred and a few thousand cells (Noble et al., 2019). We therefore consider *K* values of 64, 512 and 4,096.

Other parameter values are listed in Supplementary table 1.

### Computational model

Additional features of the computational model have been previously described (Noble et al., 2019) and the code is shared in a public repository (Noble 2019).

---

**Supplementary figure 1.**
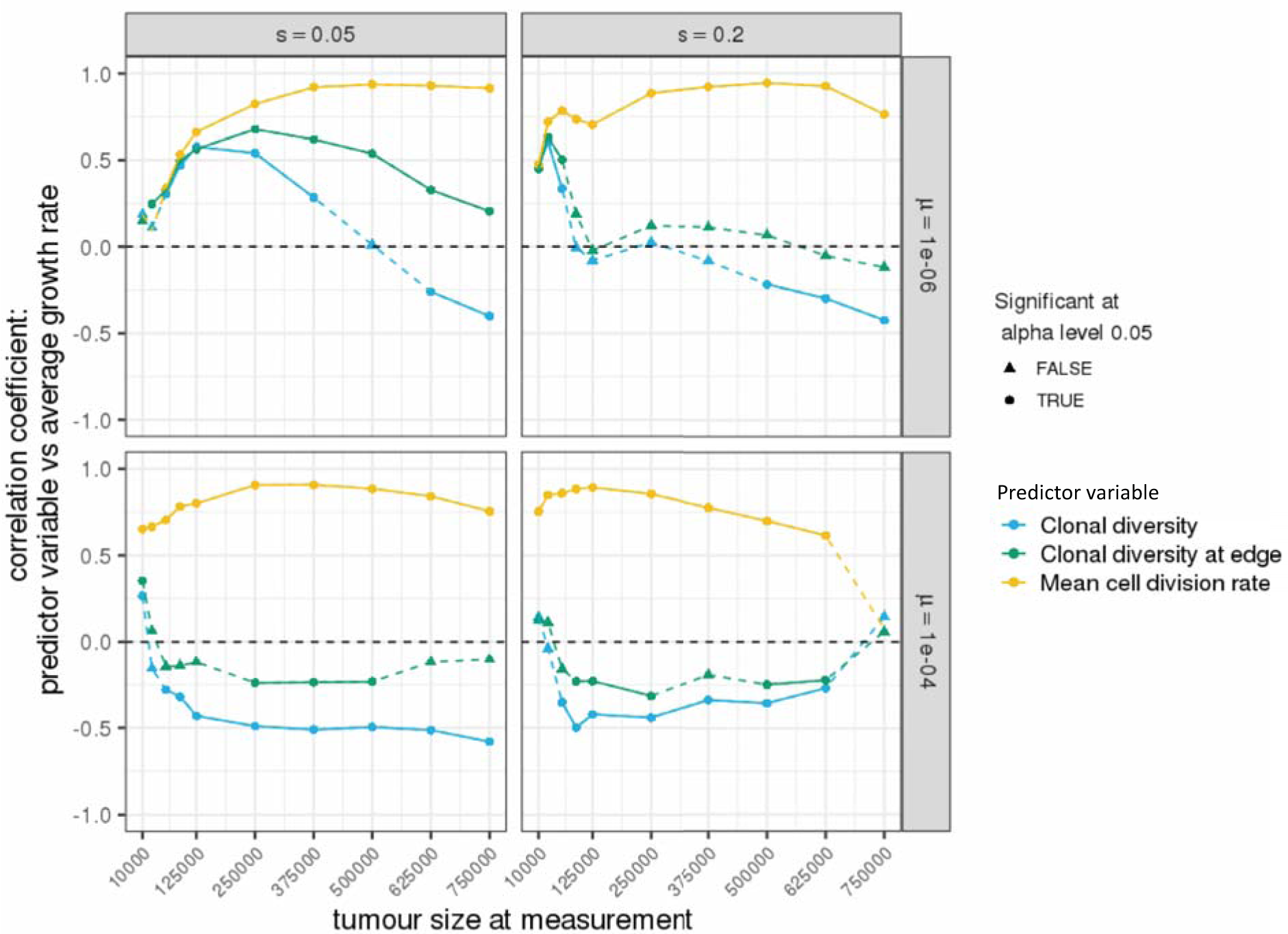
Correlation coefficients between predictor variables and future tumour growth rate, for cohorts of tumours with identical parameter values. Correlation coefficients that significantly differ from zero (p < 0.05) are indicated by circular points and solid lines, as opposed to triangular points and dashed lines. Each point represents a cohort of 100 simulated tumours; lines are added only to guide the eye.

**Supplementary figure 2.**
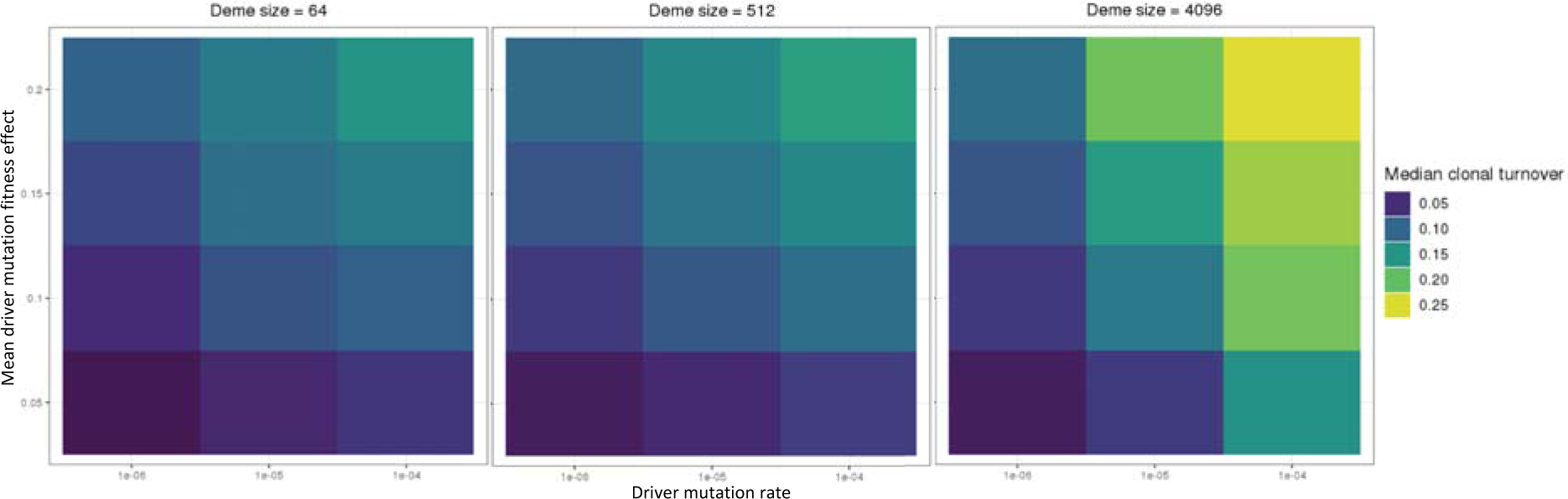
Association between parameter values and clonal turnover index. Each heatmap square is the median value of a cohort of 100 simulated tumours. Varied parameters are deme carrying capacity (*K*), driver mutation rate (*μ*) and driver fitness effect (*s*).

**Supplementary figure 3.**
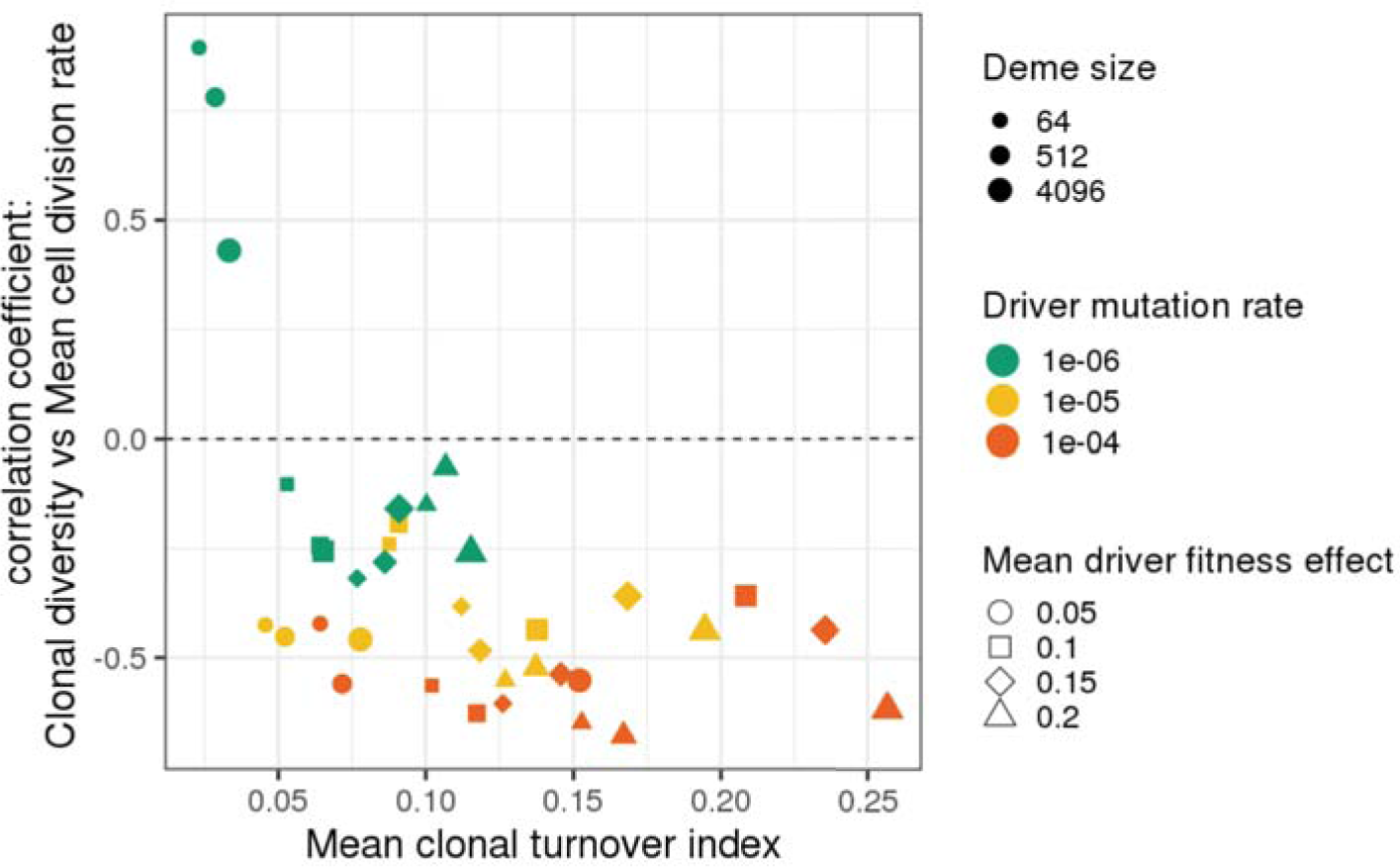
Higher clonal diversity can be associated with lower tumour cell division rate. Correlation coefficients between clonal diversity (measured at a tumour size of 250,000 cells) and mean cell division rate are plotted against mean clonal turnover index. Each point represents a cohort of 100 simulated tumours.

**Supplementary figure 4.**
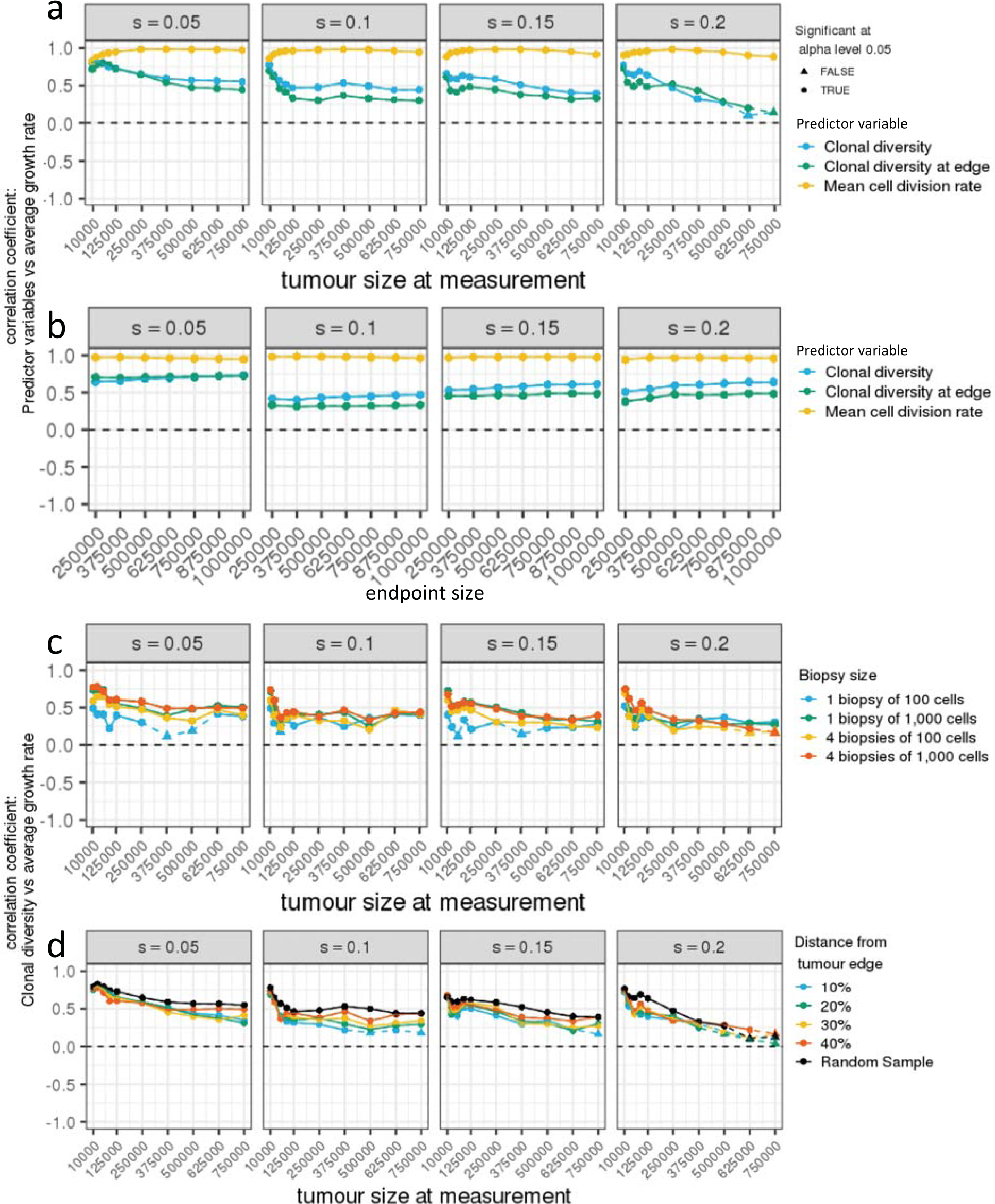
Correlations coefficients for tumours with diverse driver mutation rates. a) Correlation coefficients between predictor variables and subsequent tumour growth rate, for different measurement sizes. b) Correlation coefficients between predictor variables (measured at a tumour size of 125,000 cells) and subsequent tumour growth rate, for different endpoint sizes. c) Correlation coefficients between future tumour growth rate and clonal diversity, measured in biopsy samples of different number and size (at a depth of 40% from the tumour edge), for different measurement sizes. d) Correlation coefficients between future tumour growth rate and clonal diversity, measured in biopsy samples taken at different depths relative to the tumour edge (four biopsy samples, each of 1,000 cells, are taken from each tumour), for different measurement sizes. Correlation coefficients that significantly differ from zero (p < 0.05) are indicated by circular points and solid lines, as opposed to triangular points and dashed lines. Each point represents a cohort of 100 simulated tumours; lines are added only to guide the eye. Deme carrying capacity *K* = 512. The driver mutation rate of each tumour is chosen at random by sampling a value *X* from a uniform distribution between 4 and 6 and then setting *μ* = 10^−*X*^.

**Supplementary figure 5.**
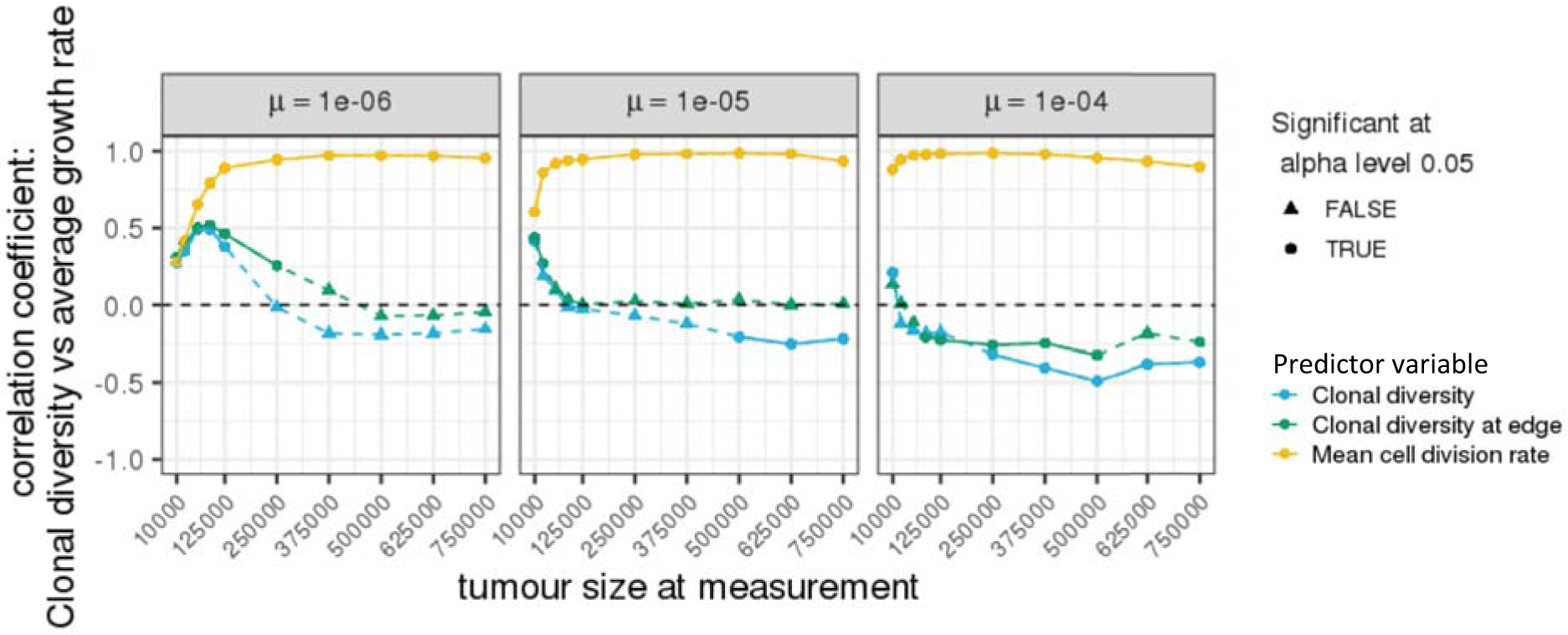
Correlations between predictor variables and future tumour growth rate, for cohorts of tumours with diverse driver fitness effects. Correlation coefficients that significantly differ from zero (p < 0.05) are indicated by circular points and solid lines, as opposed to triangular points and dashed lines. Each point represents a cohort of 100 simulated tumours; lines are added only to guide the eye. Deme carrying capacity *K* = 512. The mean driver fitness effect for each tumour is chosen at random by sampling from a uniform distribution between 0.05 and 0.2.

**Supplementary figure 6.**
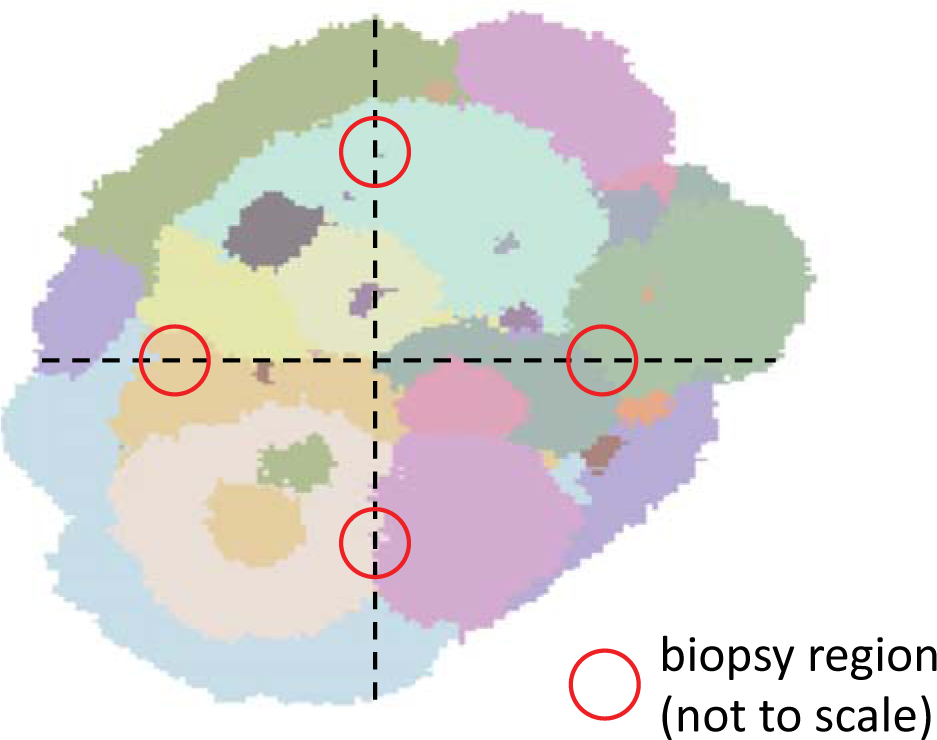
Biopsy sampling method. The centre of each disc-shaped biopsy region is located at a specified proportional depth between the tumour’s edge and its centre of mass. In this example, four biopsies are taken at a depth of 40%.

**Supplementary table 1.**
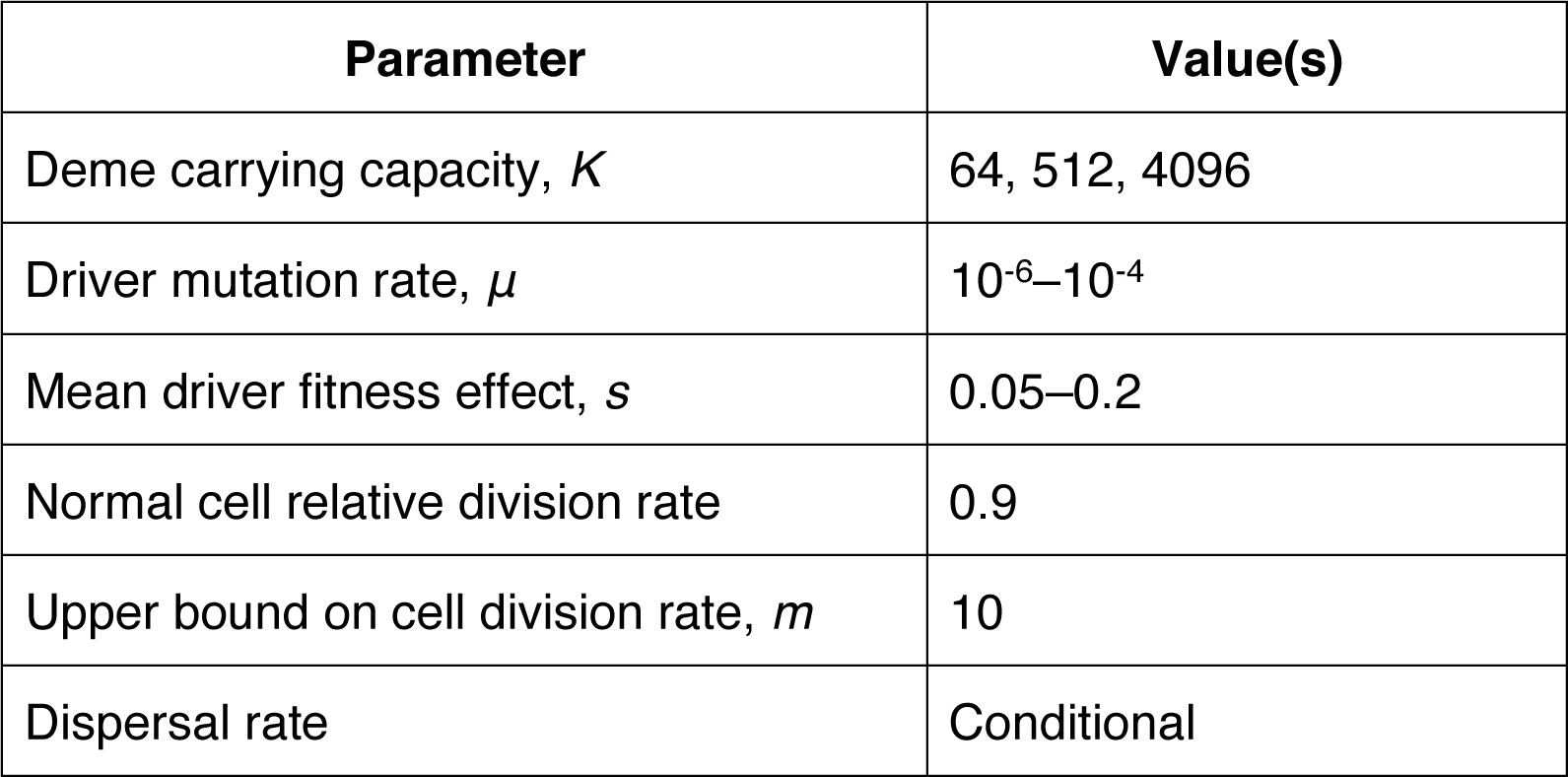
Parameter values used in this study. Mutation rate is measured per cell division; division rate is measured relative to the division rate of the initial tumour cell. The effect of a driver mutation with effect size *s* is to multiply the cell division rate *r* by a factor of 1 + *s*(1 – *r* / *m*), where *m* is the upper bound. Dispersal rates are set such that tumours typically take between 500 and 1,000 cell generations to grow from one to one million cells, corresponding to several years of human tumour growth.

| Parameter | Value(s) |
| --- | --- |
| Deme carrying capacity, $K$ | 64, 512, 4096 |
| Driver mutation rate, $\mu$ | $10^{-6}$–$10^{-4}$ |
| Mean driver fitness effect, $s$ | 0.05–0.2 |
| Normal cell relative division rate | 0.9 |
| Upper bound on cell division rate, $m$ | 10 |
| Dispersal rate | Conditional |

